# Operating CRISPR/Cas12a in a complex nucleic acid sequence background

**DOI:** 10.1101/2025.07.26.666951

**Authors:** Henning Hellmer, Thomas Mayer, Friedrich C. Simmel

## Abstract

Since their discovery, CRISPR–Cas systems have been widely applied in areas ranging from genome editing to biosensing, owing to their specific, RNA-guided target recognition. Their performance in complex biological environments has been extensively studied, particularly to optimize guide RNA design and minimize off-target cleavage. Here, we focus on the kinetic inhibition of the interaction between Cas12a - a Class 2, Type V effector - and its target, caused by interference from non-cognate background nucleic acids. This effect is particularly relevant for sensing applications in complex mixtures or cellular contexts, where genome- and transcriptome-derived sequences may impede CRISPR–Cas activity. Using in vitro assays under defined conditions, we systematically examine the influence of background single-stranded RNA (ssRNA) and double-stranded DNA (dsDNA) on reaction kinetics. We find that both the purine-to-pyrimidine ratio and the GC content of the guide RNA seed region significantly affect kinetic inhibition by background polynucleotides. Guide RNAs with low GC content and a high purine fraction in the seed region were least affected by background sequences. Experiments with dCas12a-based gene activation in living cells indicate that our in vitro findings may also be relevant for in vivo applications.

## INTRODUCTION

In recent years CRISPR/Cas technology has become a standard approach for the manipulation of genomic DNA. A wide range of CRISPR-associated (Cas) proteins have been discovered, which carry out diverse functions by binding to and chemically processing DNA or RNA molecules, or both (1). Among the most popular proteins are the CRISPR-associated endonucleases Cas9 and Cas12a, which have already found widespread use in genome editing and a plethora of other applications (2,3). The reason for the huge impact of CRISPR-associated nucleases is the fact that they are directed towards their target sequences by an RNA guide (gRNA), which may be a single-stranded RNA or a complex of several RNAs (2,3). This makes Cas proteins, in a sense, “sequence-programmable” (4). Upon incorporation of the gRNA into the Cas endonuclease, the resulting ribonucleoprotein complex scans potential target nucleic acids for complementary sequences, and after target recognition cleaves them. Besides sequence-specific cutting of the target, some RNA-guided Cas endonucleases show additional unspecific cleavage of non-cognate nucleic acids after target recognition. In its biological context this has been interpreted as a suicide reaction towards phage infection (5,6).

In the present work we focus on Cas12a, a Class 2 Type V nuclease which targets dsDNA and cleaves the double strand at the binding site (cis-cleavage) (3). Compared to the more widely used Cas9, Cas12a employs shorter, single-stranded gRNAs. In addition, Cas12a can process precursor RNA strands that contain its cognate handle sequence, thereby trimming longer gRNA transcripts into functional guides (3). Binding to its target sequence triggers a conformational change which activates the Cas protein’s cleavage domain (7). Activated Cas12a also exhibits unspecific “trans”-cleavage of ssDNA in the proximity of its nuclease domain, which can be utilized as a readout for target recognition and activation. While cis-cleavage is a single turnover process, trans-cleavage occurs multiple times, with a reported turnover rate of 17 s^-1^ (5). Short single-stranded DNA (ssDNA) reporter molecules, labeled with a fluorophore at one end and a quencher at the other, are cleaved by Cas12a’s trans-cleavage activity. This cleavage separates the fluorophore from the quencher, resulting in an increase in fluorescence that serves as a detectable signal - a readout technique that has been coined DETECTR (5,8).

Because of its unique potential for precise genetic manipulation, CRISPR/Cas technology holds significant promise in biomedicine, with genome editing emerging as one of its most far-reaching applications (9). In such settings, however, it is essential that genomic edits are both predictable and reliable within the complex biological environment of a living organism. One important and extensively studied aspect of CRISPR/Cas systems is therefore their specificity in targeting only the intended genomic locus. To achieve target precision, engineered Cas variants with enhanced binding fidelity have been developed, along with AI-guided tools that predict potential off-target sites across the genome (10,11).

While off-target effects and cleavage kinetics have been studied extensively, the specific impact of competing RNA molecules and non-target DNA on target cleavage kinetics has received comparatively less attention. CRISPR/Cas target recognition in cells involves a series of steps that must occur before DNA cleavage can proceed (12). First, the expressed Cas protein must bind the transcribed gRNA, forming a functional Cas–gRNA complex. This complex then searches the genome for its target sequence (13,14). Two distinct types of interactions may influence the kinetics of these early steps. During and shortly after transcription, unbound gRNA may interact with other RNAs in the nucleus, potentially leading to sequestration or the formation of secondary structures that interfere with Cas protein binding (15). Once the gRNA is successfully incorporated into the Cas protein, the target search begins. In this phase, the Cas–gRNA complex scans the DNA for a protospacer adjacent motif (PAM) – a short, sequence motif essential for initial binding and local DNA unwinding. For Cas12a, the PAM sequence is 5’-TTTV-3’, where V represents A, C, or G. Upon PAM recognition, the complex interrogates the adjacent DNA sequence for complementarity to the gRNA spacer. When sufficient complementarity is not found, the complex dissociates and resumes scanning. The first eight nucleotides (nt) adjacent to the PAM – the so-called seed region – have been found to be particularly critical for initiating hybridization. When sufficient complementarity is established, the complex undergoes a conformational transition into the R-loop state, in which the non-target DNA strand (sharing the same sequence as the gRNA spacer) is displaced. This structural rearrangement positions the nuclease domain for cleavage, enabling site-specific cutting of the target DNA. In Cas12a, this activation also triggers trans-cleavage activity (16–18).

The stability of the seed region plays a critical role in determining the dissociation rate of the Cas-gRNA-DNA complex. This implies that the sequence composition of the gRNA, and consequently the target, influences the duration for which the Cas-gRNA remains bound at an off-target DNA locus containing a PAM sequence, where complementarity is probed. Stronger interactions between gRNA and off-target DNA are expected to result in prolonged binding, slowing down the kinetics of the intended reaction. It has in fact been shown that CRISPR-Cas off-target binding can be well understood in the context of nucleic acid hybridization kinetics (19).

Using single molecule tracking experiments (13,14,20,21), the kinetics of the target search process have been investigated for the Cas9 protein in greater detail. The search process is a combination of three-dimensional diffusion between DNA strands or strand regions and one-dimensional diffusion along the DNA once the Cas9 binds to a PAM sequence. Dwell times at a PAM site are typically on the order of one second but increase substantially with more PAMs or target sequences adjacent to the initial binding site (21,22). Other studies revealed that interactions between the Cas12a effector and DNA are similar to those for Cas9 (23,24).

It has previously been noted that the recognition and binding of CRISPR-associated effectors to their target sites, which requires both a PAM and a complementary sequence, resembles a toehold-mediated strand displacement (TMSD) process (25,26). TMSD is a widely employed mechanism in nucleic acid nanotechnology, where an invading strand displaces an incumbent strand from a nucleic acid duplex through a branch migration process. Recent studies have shown that TMSD processes are impacted by the sequence composition of the participating nucleic acid strands in various ways. For instance, the sequence of the invader strand dictates its interactions with background sequences, which slow down the binding of the invader to the target (27). Moreover, once strand displacement is initiated, it has been found that individual steps of the branch migration process are highly affected by the sequence composition, particularly in RNA–DNA hybrid strand displacement processes (28,29).

In the present study, we investigate how the sequence composition of gRNAs influences the kinetics of the intended Cas–gRNA–target interaction in vitro within complex sequence environments. Specifically, we employed synthetic pools of background double-stranded DNA (dsDNA) containing a variety of sequence motifs. These include sequences with partial complementarity to the gRNA but lacking a PAM site, sequences with complementarity only in regions distal to the seed, and other combinatorial configurations. We further analyzed the effects of background pools consisting of random dsDNA or ssRNA, each pool potentially covering the full spectrum of possible gRNA target motifs.

To further explore the impact of gRNA composition, we also employed sequences based on different three-letter alphabets and varied both the purine-to-pyrimidine ratio and GC content. This enabled us to probe sequence design principles that were previously shown to influence DNA/RNA hybridization and branch migration kinetics in the context of TMSD. Based on the quantitative analysis of our data, we identified sequence features that correlate with reduced kinetic inhibition. We hypothesize that the found sequence features mitigate off-target interactions and unfavorable secondary structures, thereby facilitating more efficient target recognition. We finally examined a CRISPR/Cas system designed based on these features also in living HEK cells. Our results suggest that in the complex intracellular environment, careful selection of target site sequences can indeed enhance the kinetics of CRISPR/Cas-mediated activity.

## MATERIALS AND METHODS

### Materials

We used NUPACK for all sequence design and analysis tasks (30,31). Template oligos for gRNA transcription were ordered as PAGE-purified dsDNA at Integrated DNA Technologies (IDT), targets were ordered as lab-ready dsDNA in 1x IDTE buffer. Templates and primers for the preparation of random ssRNA background and dsDNA background were ordered as lab-ready ssDNA in 1x IDTE buffer. Cas12a (Alt-R™ AsCas12a Ultra nuclease) was ordered from IDT, reaction buffer (NEBuffer 3.1) was ordered from New England Biolabs (NEB). Single-stranded, fluorophore-quencher labeled reporter strands (5’-FAM-TTATT-BHQ1-3’) were ordered from IDT. Q5 mastermix for PCR reactions as well as in vitro transcription (IVT) components, namely T7 RNA polymerase, rNTPs and RNA polymerase buffer were ordered from NEB. For DNA digestion we used TURBO DNA-free™ kits from Thermo Fisher. For RNA and DNA purification, Zymo Research RNA or DNA Clean & Concentrator kits were used. Gene fragments for cloning were ordered at Twist Biosciences, short DNA oligos were ordered from IDT in 1x IDTE buffer. BsmBI, T4 ligase and T4 polynucleotide kinase were purchased from NEB. Bacterial host cells (*E. coli* DH5alpha strain) and LB medium were ordered from Thermo Fisher. For cell culture, HEK293T cells were acquired from ATCC. DMEM and Opti-MEM media, FBS, DPBS and Trypsin-EDTA were ordered from Merck. Antibiotic-antimycotic and Lipofectamine™ 2000 were purchased from Thermo Fisher.

### In vitro assay overview

All in vitro experiments were performed following the same procedure. The final assay composition contained 1x reaction buffer (NEBuffer 3.1), 60 nM AsCas12a, 150 nM gRNA, 6.7 nM dsDNA target, 200 nM ssDNA reporter and either 2 µM of RNA background strands, 200 nM of DNA background strands or no background. The mixture was brought to a final volume of 15 µl with water. A mastermix containing all constituents except for target DNA and guide RNA was prepared. The mastermix was pipetted into a Corning® low volume, non-binding plate and the gRNA and the target were added last. The plate was sealed, samples spun down and the plate was put into either a BMG Labtech FLUOstar® Omega or CLARIOstar® plate reader with the temperature set to 37°C and excitation and emission filters were preset to 485 nm (15 nm bandwidth) and 530 nm (20 nm bandwidth) respectively. After starting the measurement, fluorescence intensity data was collected each minute for 16 hours. The data was transferred to an Excel file and subsequently plotted and analyzed using custom-python scripts (provided in the GitHub repository). For every measurement a negative control without target DNA was conducted. Experiments were performed in technical triplicates on the same plate.

### Preparation of guide RNAs and polynucleotide backgrounds

#### guide RNAs

Guide RNAs (gRNAs) were transcribed from dsDNA oligos containing the T7 RNAP promoter sequence (provided in the GitHub repository). Transcription was carried out following the standard NEB protocol, with an incubation time of 16 hours to maximize RNA yield. Transcripts were treated with TURBO^TM^ DNase according to the rigorous treatment protocol, followed by purification with the RNA Clean & Concentrator kit (Zymo Research).

#### dsDNA background

The dsDNA background used in our experiments was only partially random, as we had to balance sequence randomness against synthesis yield and cost. To our knowledge, commercially available dsDNA cannot be ordered with fully randomized sequences — only ssDNA is available in this format. As a workaround, we ordered 200-nucleotide-long ssDNA oligonucleotides containing two fixed primer-binding regions at the 5′ and 3′ ends (5’-P1– N168–P2-3’), resulting in 168 central nucleotides with randomized composition. The sequences of the constant flanking regions – which were designed to not contain a PAM for Cas12a – are provided in the GitHub repository. To reduce synthesis costs, we amplified these ssDNA templates by PCR, ensuring that each resulting dsDNA molecule exists in multiple copies. Even though the presence of the fixed primer sequences limits the overall sequence diversity, there are still 168 random bases per strand, corresponding to 4^168^ ≈ 10^100^ different possibilities for the sequence. The number N of different sequences before performing the PCR can be calculated via *N* = *N*_*A*_ · *c*(*ssDNA*) · *V*, where *N*_*A*_ is the Avogadro constant, *c*(*ssDNA*) the concentration and *V* the volume of the ssDNA solution, which in our case results 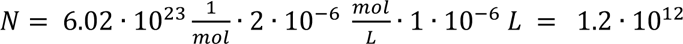 different sequences to start with. For the PCR we added the forward and reverse primer to a final concentration of 2 µM, for which we obtained the best yield of the full-length product without generating shorter dsDNA strands due to undesired primer binding.

#### ssRNA background

In the case of ssRNA background we were able to produce fully random ssRNA strands. To this end, we ordered ssDNA with the 183 random nucleotides, followed by the reverse complementary T7 promoter sequence. Before transcription we added an oligo with the T7 promoter sequence that can bind to its reverse complement and added the Q5 mastermix and performed a primer extension reaction, to produce a double stranded template for IVT. As for the gRNAs, the transcription was performed according to the standard NEB protocol with transcription time set to 16h to maximize the output yield. The IVT output RNA with a length of 183 bases was processed with the TURBO DNA-free™ Kit and subsequently cleaned up using the Zymo Research RNA Clean & Concentrator kit.

### Kinetic data analysis

Kinetic data was analyzed using Excel and python. The data is provided in a time vs. intensity format. We performed technical triplicates and included negative controls in every measurement. For normalization and better comparability, the background value (obtained from the negative controls) was subtracted from the intensity values and reactions normalized to a final value of 200 (which is the nanomolar concentration of reporter in every measurement). For comparison of the kinetics with (w) and without (w/o) background sequences we extracted the time to reach 50% completion (*t*_0.5_) for each sample with each condition by identifying the earliest time point at which the fluorescence intensity exceeded 50% of the maximum signal observed for that sample. To compare the effect of background DNA on kinetic performance we computed all possible pairwise *t*_0.5_ ratios via 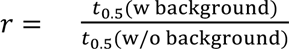 between replicates of the DNA-containing condition and its matching DNA-free control (i.e., 3×3 = 9 comparisons per group). The mean of these nine ratios was taken as the relative t₀.₅, and the standard deviation was used to represent the variability across comparisons.

### Symbolic Regression

We applied symbolic regression using the PySRRegressor from the pysr library to identify algebraic expressions that relate features of our spacer sequences to the measured kinetic slowdowns. Sequence features from the seed region that appeared to correlate with the kinetic ratio, such as the purine-to-pyrimidine ratio, were, together with the kinetic ratio, imported into a data frame for further analysis. We used the PySRRegressor, minimized the mean squared error with a custom element-wise loss function, and selected models based on the best loss score. To prevent the generation of uninterpretable expressions, we constrained model complexity by setting a maximum depth of 6 and a maximum expression size of 8. For our final model, we excluded three gRNA sequences (1, 6, and 7) from training because they skewed the resulting equation, making it uninterpretable. These gRNAs were therefore considered outliers. Our goal was to capture the overall trend rather than derive overly complex formulas. The full code and dataset are available in the accompanying GitHub repository.

### ODE model

For the model used in the discussion around Figure 6, we first fitted the data from a DETECTR experiment using gRNA 2 in the absence of a nucleic acid background. To this end, we normalized the fluorescence data to span a range corresponding to 0 to 200 nM target concentration. We used scipy.solve_ivp as the ODE solver and manually adjusted *k*_ON,T_ and *k*_2_ within a physically reasonable range until agreement with the data was found. These two parameters were then fixed. Using the dataset for gRNA 2 in the presence of a random DNA background, the remaining two rate constants of Model 2 (*k*_ON,B_, *k*_*OFF,B*_) were determined in the same way. The reported values are consistent with typical rates for such processes.

### *In cellulo* experiments

For *in cellulo* experiments the readout strategy had to be adapted. We chose a denCas12a-VPR based CRISPR activation system that results in GFP expression in response to target binding. Plasmids for the gRNA and a denCas12a-VPR construct, respectively, were transiently transfected alongside a target plasmid coding for GFP.

#### Plasmid Design and Cloning

The plasmid coding for constitutive expression of denAsCas12a-VPR was available in-house. The gRNA plasmids were cloned via Golden Gate assembly (32). As backbone we used a plasmid with the required cutting sites (BsmBI) downstream of a basic U6 promoter. An additional U6+27 sequence and the gRNA sequence were designed in Benchling and ordered as single-stranded oligonucleotides with 4-base overlaps. After hybridization, the two resulting duplexes and the backbone were joined together in an assembly reaction following standard procedures and supplemented with T4 poly-nucleotide kinase to phosphorylate the ordered DNA strands. For the target plasmid longer gene fragments were designed. These included a CRISPR target site for the respective gRNA sequence followed by a minimal promoter in front of a green fluorescent protein (GFP) gene. The designed DNA duplexes were appended on each side with BsmBI cutting sites. As backbone, a part of a plasmid was copied via PCR introducing complement BsmBI cutting sites. This part introduced a bovine growth hormone polyadenylation (*bgh*-PolyA) signal after the GFP gene upon correct assembly. All plasmid sequences are listed in the GitHub repository.

#### Cell Culture and Experimental Procedure

The performance of CRISPR activation with different gRNAs in a cellular environment was tested in HEK293T cells, which were cultured under standard conditions. The cells were seeded onto a 48-well plate at 50.000 cells per well and transfected with the three plasmids (target, gRNA and denCas12a-VPR plasmids) 24 hours later. After 10 hours of incubation after the transfection, the plate was put on the stage of an EVOS M7000 microscope with an incubation chamber setup. Brightfield and GFP channel images were then taken for 17 h in 10-minute intervals. Image analysis was conducted with Fiji and Python (script available in the GitHub repository). The images were stacked and subdivided into a 30-by-30 grid, with each grid segment approximately matching the size of the HEK293T cells. The intensity of GFP fluorescence of each resulting image stack segment was plotted against time. The image background was corrected with manually selected image regions that did not exhibit any fluorescence. Regions that did not show an increase of fluorescence over time were omitted from further analysis, while the remaining intensity curves were summed up and divided by their number. The resulting average fluorescence increase was fitted with a linear function. Different conditions (gRNA/target sequences) were further compared by the averaged fluorescence intensity levels after 17h. The experiments was conducted in biological triplicates (three seeded wells per condition). Image series were acquired from two fields of view per well.

## RESULTS

### Impact of designed background strands on interaction kinetics

The Cas/gRNA target search process has been previously studied in great detail, which has led to a good understanding how this ribonucleoprotein complex unwinds dsDNA in an RNA-DNA hybrid strand displacement process, during which gRNA displaces the non-target strand and binds to its target sequence.

It has been shown that the first few bases after the PAM recognition site, which are called the seed region, are the most important for stable, initial binding of the Cas-gRNA complex to the dsDNA (3,7). For the Cas12a protein, the first 5-6 bases downstream of the PAM are typically regarded as the seed sequence (7,18). However, mismatches within the first ≈ 10 bases are known to significantly impair target recognition and binding (33). For our experiments and designs, we thus regarded the first 8 bases after the PAM as the relevant seed sequence (Fig. 1A). To assess how different Cas12a/gRNA combinations perform in the presence of a background composed of designed or random nucleic acid sequences, we used the DETECTR method for readout (Fig. 1B, cf. Methods).

**Figure 1.**
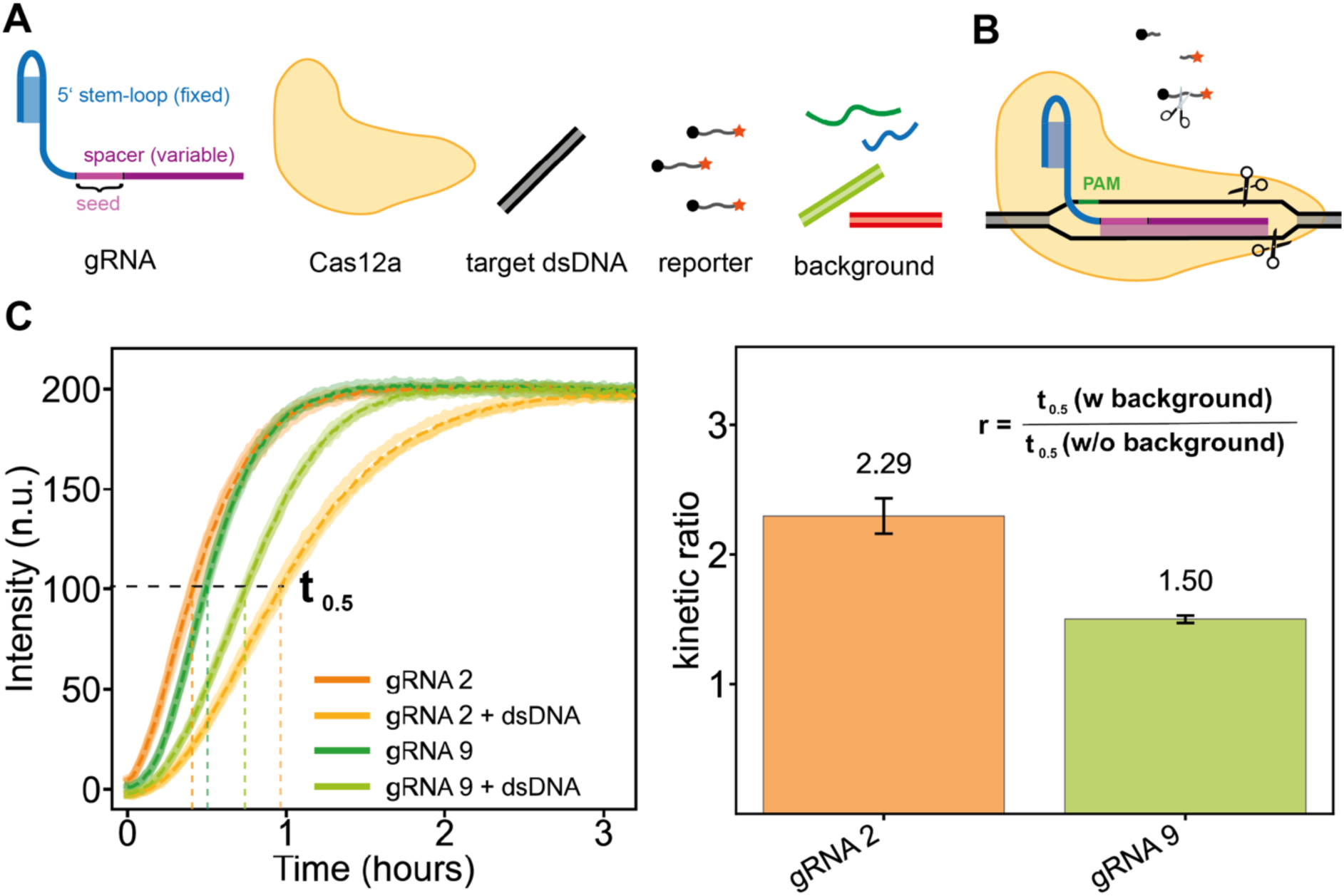
Experimental in vitro setup and analysis. (**A**) Individual components of the experiment. The gRNA contains a fixed region required for recognition and uptake by the Cas12a protein. The downstream variable region is called the spacer, with the first eight bases referred to as the seed region. Additionally, we used target dsDNA, a ssDNA reporter labeled with a fluorophore–quencher pair, and varying backgrounds of dsDNA and ssRNA. (**B**) Trans-cleavage property. Upon successful binding to the target dsDNA, Cas12a exhibits trans-cleavage activity, meaning it non-specifically cleaves surrounding ssDNA. This property can be used to measure system activity in a plate reader experiment. (**C**) Data acquisition and analysis. **Left:** Fluorescence curves measured for two different gRNAs, with and without dsDNA background. Although gRNA 2 shows the fastest kinetics without dsDNA background, gRNA 9 is faster in the presence of a dsDNA background. **Right:** Comparison of the t₀.₅ values as a measure of the slowdown caused by the background. The gRNA 2 system is slowed down more (r = 2.32) than the gRNA 9 system (r = 1.50), indicating that gRNA 9 is less affected by the dsDNA background.

Upon binding to its target and activation of its nuclease function, Cas12a exhibits a collateral trans-cleavage activity on nearby DNA strands. When fluorogenic probes - labeled with both a fluorophore and a quencher - are supplied, this collateral activity leads to a fluorescence increase, which serves as a proxy for successful target recognition and activation. As illustrated in Fig. 1C, DETECTR can be employed as a readout to compare the performance of different guide RNA types in the presence or absence of interfering background strands. As demonstrated by the representative fluorescence traces shown, different gRNA types are affected by the background to varying degrees. In the following, we quantify the reaction kinetics by determining the half-time t_0.5_ for completion of each process. The effect of the background is then expressed as the ratio r of the t_0.5_ values in the presence and absence of background strands, respectively.

To study kinetic inhibition by non-cognate DNA sequences that can hinder the target search process, we designed four different versions (V1-V4) of a dsDNA background. V1 contained the PAM sequence, no complementarity for the seed region of the gRNA but perfect complementarity for the remaining 12 bases. V2 lacked the PAM but contained a sequence perfectly matching the gRNA spacer, V3 harbors a matching PAM and target for the seed sequence but no complementarity in the remaining part. Lastly, V4 has multiple PAMs, but is not complementary to the spacer at all. As explained in Fig. 2A, we expected these different types of backgrounds to interfere at different stages of the search process.

**Figure 2.**
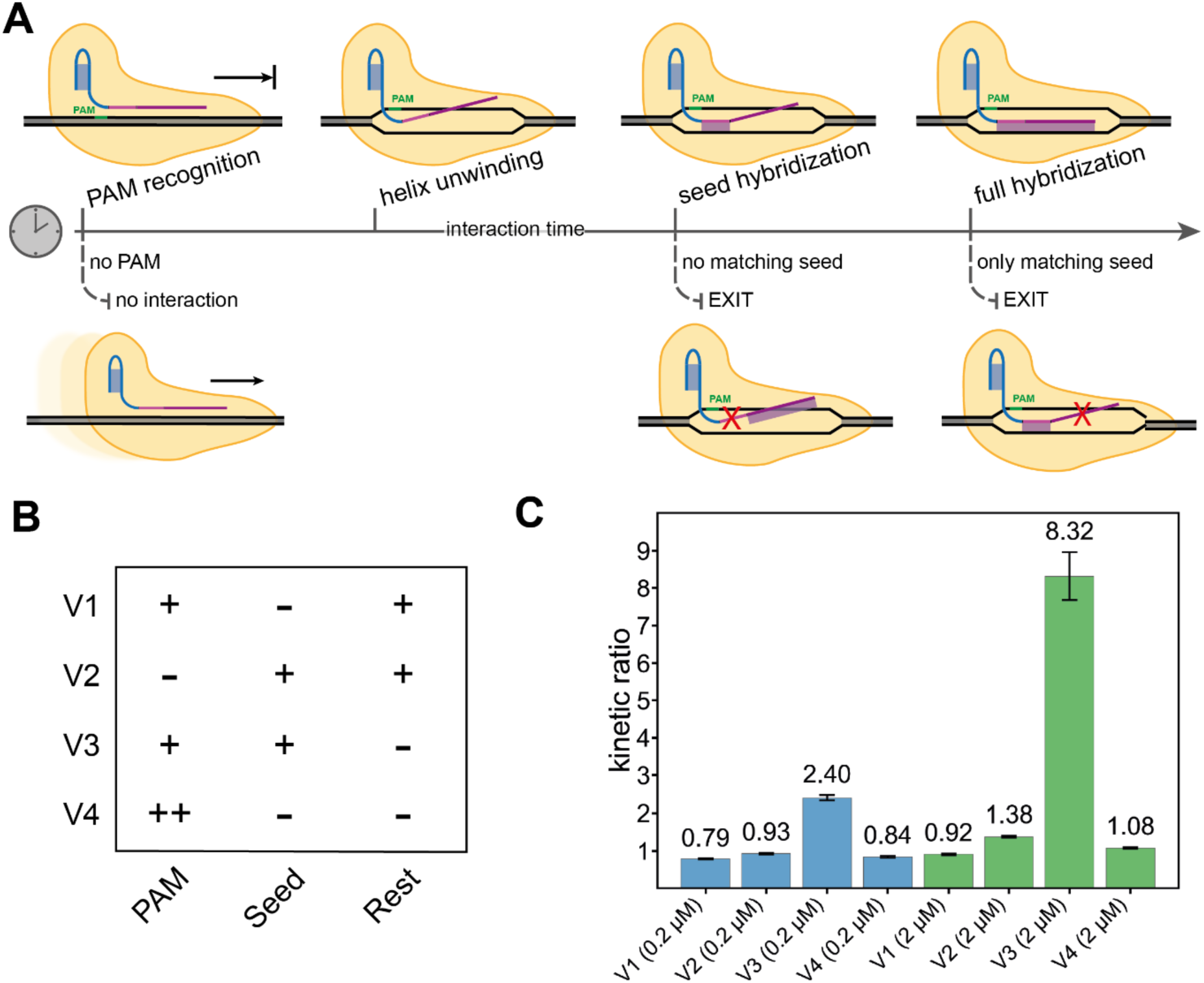
Cas12a target recognition. (**A**) Modes of interaction between the Cas12a enzyme and background DNA and the corresponding interaction time for any DNA locus. Cas12a loaded with a guideRNA is moving along the DNA scanning for protospacer adjacent motifs (PAMs). Upon binding to a PAM the DNA helix is unwound, and the spacer hybridization takes place starting at the seed region. Only if the complete sequence matches the DNA, R-loop formation is completed, and the endonuclease activity is unlocked. (**B**) Sequence designs for background DNA harboring parts of an additional target sequence to compete with the original target in the assay. V1 carries a PAM and the target sequence missing the seed region, V2 incorporates the complete target but lacks a PAM, V3 only displays a PAM and the seed region, V4 has several PAMs but no target sequence. (**C**) Relative t₀.₅ comparison as a measure of the slowdown caused by the specifically designed DNA background. Blue: background DNA at 0.2 µM, green: background DNA at 2 µM. The greatest slowdown is caused by background version 3 which is in line with the modes of interaction (see A) as present PAM and seed region cause the longest interaction with the background DNA.

Measurements of the kinetic ratio r (defined in Fig. 1C) for the different backgrounds are shown in Fig. 2C. A background of double-stranded DNA lacking a PAM sequence (V2) does not appear to inhibit target search by the Cas12a:gRNA complex at all, even though V2 contains a perfectly matching seed region. Moreover, we do not observe any kinetic inhibition by background V1, which includes a PAM, but no sequence matching the seed. This suggests that the Cas12a:gRNA only briefly interacts with PAM sites when the initial bases of the seed region cannot hybridize with the DNA target. For Cas9, similar interactions with isolated PAM sites have previously been shown to last approximately 1 second (21).

Even in the case of the V4 background sequences, which contain eight PAM sites each, we do not observe any kinetic inhibition within the tested concentration range. This appears to differ from previous observations made for Cas9, suggesting an even faster dissociation of Cas12a from a PAM in the absence of a matching seed region (20).

Notably, a pronounced kinetic slowdown by a factor of ≈ 8 is observed in the case of background V3 (at 2 µM concentration), which contains both a matching PAM and a seed region. Upon encountering the PAM, the Cas12a:gRNA complex can transiently bind to the first eight bases of the double-stranded DNA, reducing the unbinding rate. However, because the remaining 12 bases do not match, this partial interaction is insufficient to initiate R-loop formation and the associated conformational changes required for the activation of cis- and trans-cleavage (18).

### Impact of random dsDNA and ssRNA background strands on interaction kinetics

We next investigated the influence of dsDNA and ssRNA background strands with randomized sequences on Cas12a–gRNA target interactions. As described in the Materials and Methods, members of the dsDNA pool contain a 168 bp long randomized sequence domain, which is flanked by constant primer-binding sites required for amplification. The primer sequences were specifically designed to exclude PAM motifs and to not exhibit extensive complementarity to the gRNAs. The primer regions were therefore not expected to significantly influence the experimental outcome.

We worked with randomized dsDNA pools at a total concentration of 200 nM, corresponding to a 30-fold excess relative to the target concentration (6.7 nM). In contrast, for ssRNA we used a concentration of 2 µM, as little to no inhibitory effect was observed at lower concentrations. Due to the substantially weaker kinetic inhibition exerted by ssRNA compared to dsDNA at equivalent concentrations, we hypothesize that ssRNA influences the kinetics mainly by interacting with the gRNA prior to its incorporation into the Cas protein, but not with gRNA sequestered within a Cas12a complex. Such interactions can occur when all components are mixed simultaneously, potentially inhibiting the association of Cas12a with the gRNA.

As shown in Fig. 3B, we tested different types of gRNAs with varying sequence features (listed in Table 1) to explore how their activity is affected by the presence of random nucleic acid backgrounds. With these gRNAs, we evaluated design strategies that had previously been applied to enhance the robustness and predictability of nucleic acid strand displacement reactions (27). These included gRNA sequences based on restricted three-letter alphabets (3L-A (gRNA 3), 3L-C (gRNA 4), 3L-G (gRNA 5), 3L-T (gRNA 6)) and sequences with varying purine-to-pyrimidine ratios. Additionally, we tested a gRNA (gRNA 1) that had been used previously in our lab (34), along with a version in which its bases were randomly permuted (gRNA 2). Although secondary structure of the gRNA may also influence kinetics, we did not specifically account for it in this study. Instead, the gRNA sequences were designed to avoid strong secondary structures within the spacer sequence, which is a commonly applied design principle (35,36).

**Figure 3.**
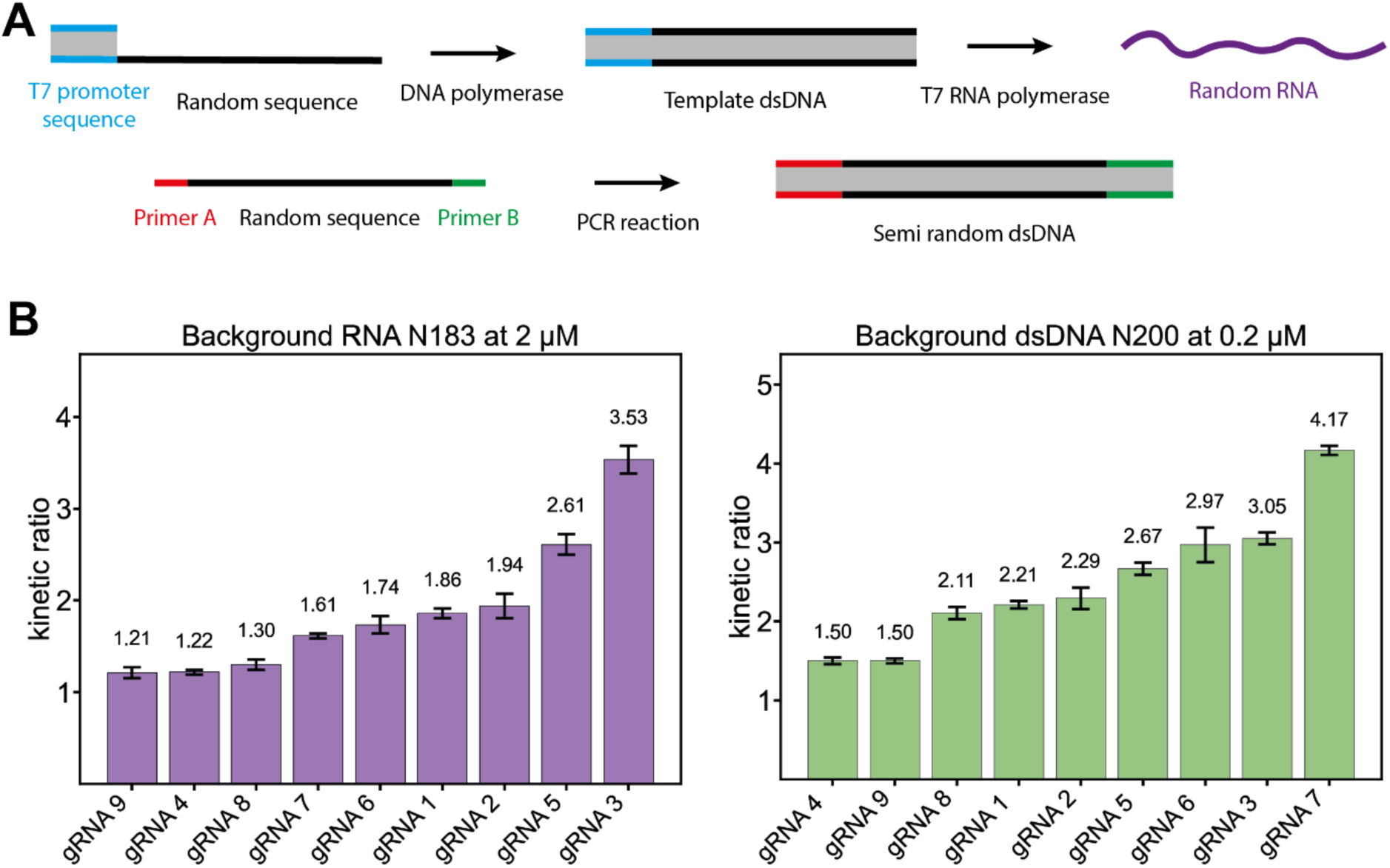
Multiple gRNAs operated in ssRNA or dsDNA background. (**A**) Production of random RNA and semi random dsDNA. A randomized DNA template with only the T7 promoter sequence fixed is used to first produce the template dsDNA and later transcribe it to random RNA. For DNA, a semi random ssDNA with two primer sequences flanking the random part is used and multiplied via a PCR reaction. (**B**) First set of gRNAs with RNA or dsDNA background. The dsDNA background (right side) showed comparable slowdown with only a tenth of the concentration of the RNA. A direct comparison between the slowdown caused by RNA and dsDNA shows that there are some gRNAS that are neither heavily influenced by the RNA nor by the dsDNA (e.g. gRNA 9) and others like gRNA 3 which are slowed down by RNA as well as dsDNA background. Another visual trend is the dependence on the purine to pyrimidine ratio, especially for the dsDNA background. The gRNA with a higher purine ratio in the seed region is less influenced by the background.

**Table 1:**
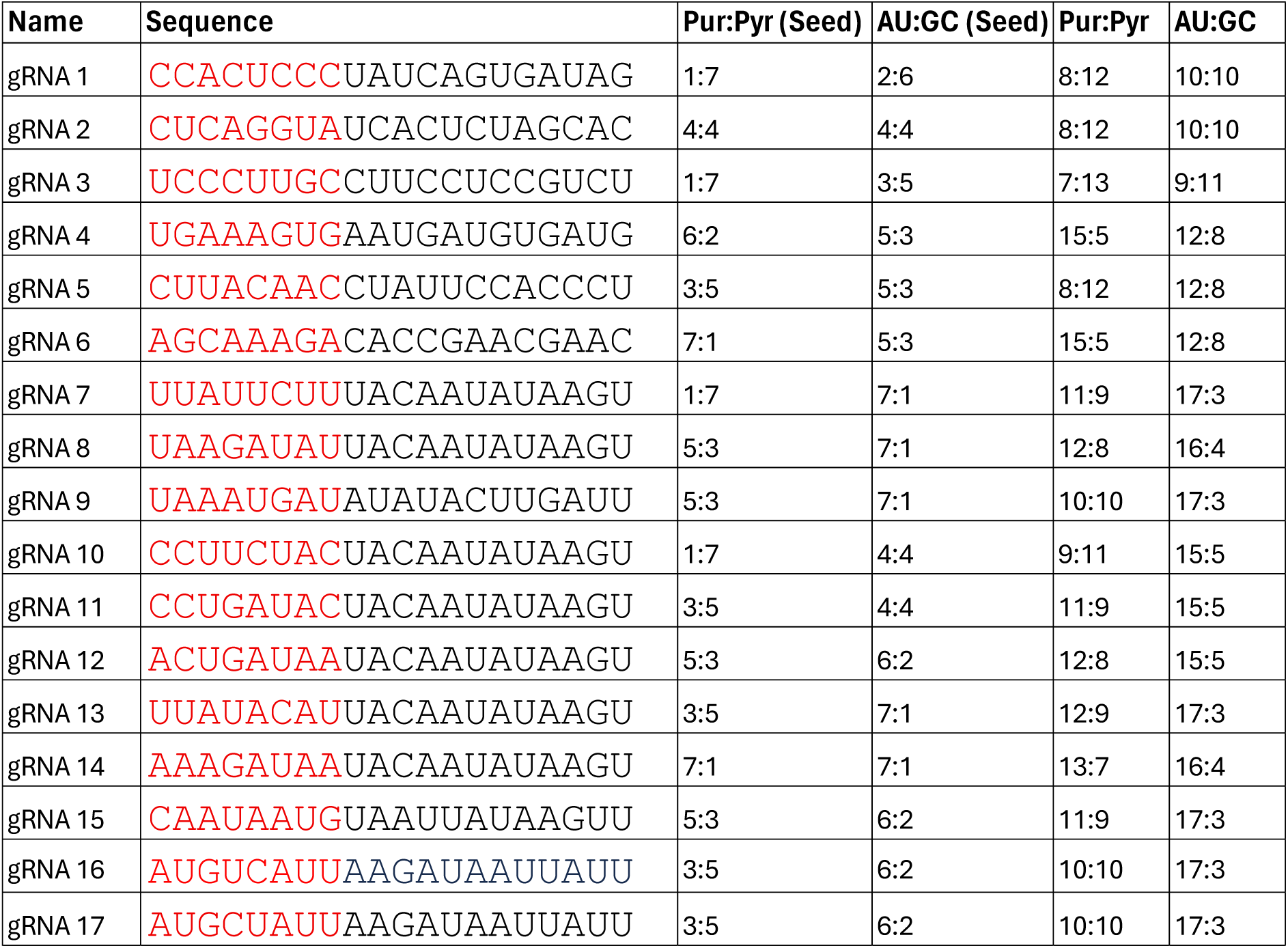
Guide RNA spacer sequences used in the experiments.

Notably, even in the absence of background strands, target cleavage kinetics assessed via DETECTR already showed substantial variation across the different gRNAs, consistent with previous observations (35,37). To allow for a meaningful comparison of kinetic inhibition across different conditions, we therefore normalized the cleavage rates by defining a kinetic ratio 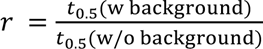 as above. Our comparison of kinetic ratios in Fig. 3B thus does not focus on the fastest kinetics, but rather on the robustness of each gRNA design in the presence of randomized nucleic acid backgrounds. The data show that the ssRNA background causes less kinetic inhibition than the dsDNA pool, although the effect still depends on the specific gRNA sequence used. Notably, the dsDNA background exerts a stronger inhibitory effect despite being present at only 10% of the RNA concentration. Some gRNAs experience strong inhibition by both dsDNA and RNA (e.g., gRNA 3), while others are largely unaffected by either (e.g., gRNA 9), and some are specifically inhibited by dsDNA alone (e.g., gRNA 7). Interestingly, our data suggest that both the purine-to-pyrimidine ratio and the GC content play a critical role in determining the extent of kinetic inhibition. To further investigate these effects, we designed a new set of gRNAs (gRNA 10-gRNA 13) with varying purine-to-pyrimidine ratios specifically in the seed region. These included sequences with either variable GC content or constant GC content to disentangle the individual and combined contributions of these two parameters. In addition, we generated further variants with low GC content (gRNA 14-gRNA 17), including a scrambled version (gRNA 16) of the original gRNA 9 to test whether the reduced inhibition observed was sequence-specific or might be a general phenomenon (cf. Table 1).

Given the stronger inhibitory effect observed with the dsDNA background, we focused our experiments on the random dsDNA pool. Using the new set of gRNAs, we repeated our analysis, as shown in Fig. 4. The results indicate a correlation between the purine-to-pyrimidine ratio and the extent of kinetic inhibition: gRNAs with higher purine content are slowed down less in the presence of the dsDNA background. Furthermore, the sequence composition of the seed appears to play a role. In order to capture these trends in a simple mathematical expression we performed a symbolic regression analysis of the kinetic ratio as a function of sequence features such as those shown in Table 1. The expression shown in the inset of Fig. 4C indeed yields lower r values for sequences with a higher purine:pyrimidine ratio in the seed region. Additionally, there is a weaker dependence on the number of cytosine bases in the seed, with larger numbers of C’s leading to increased r values. Three of the tested sequences (gRNA 1, gRNA 6, and gRNA 7) are not well described by our simple formula— most notably gRNA 7, which is the most strongly affected by the random pool. gRNA 7 has a very low purine:pyrimidine ratio of 1:7 in the seed. Compared to the other two gRNAs with the same ratio (gRNA 1 and gRNA 4), it has the highest AU:GC ratio, which in this context implies a large number of uracils. While gRNA 1 also has a low purine:pyrimidine ratio, the third outlier, gRNA 6, does not fit the trend: it has a very high purine:pyrimidine ratio and contains many adenines in the seed.

**Figure 4.**
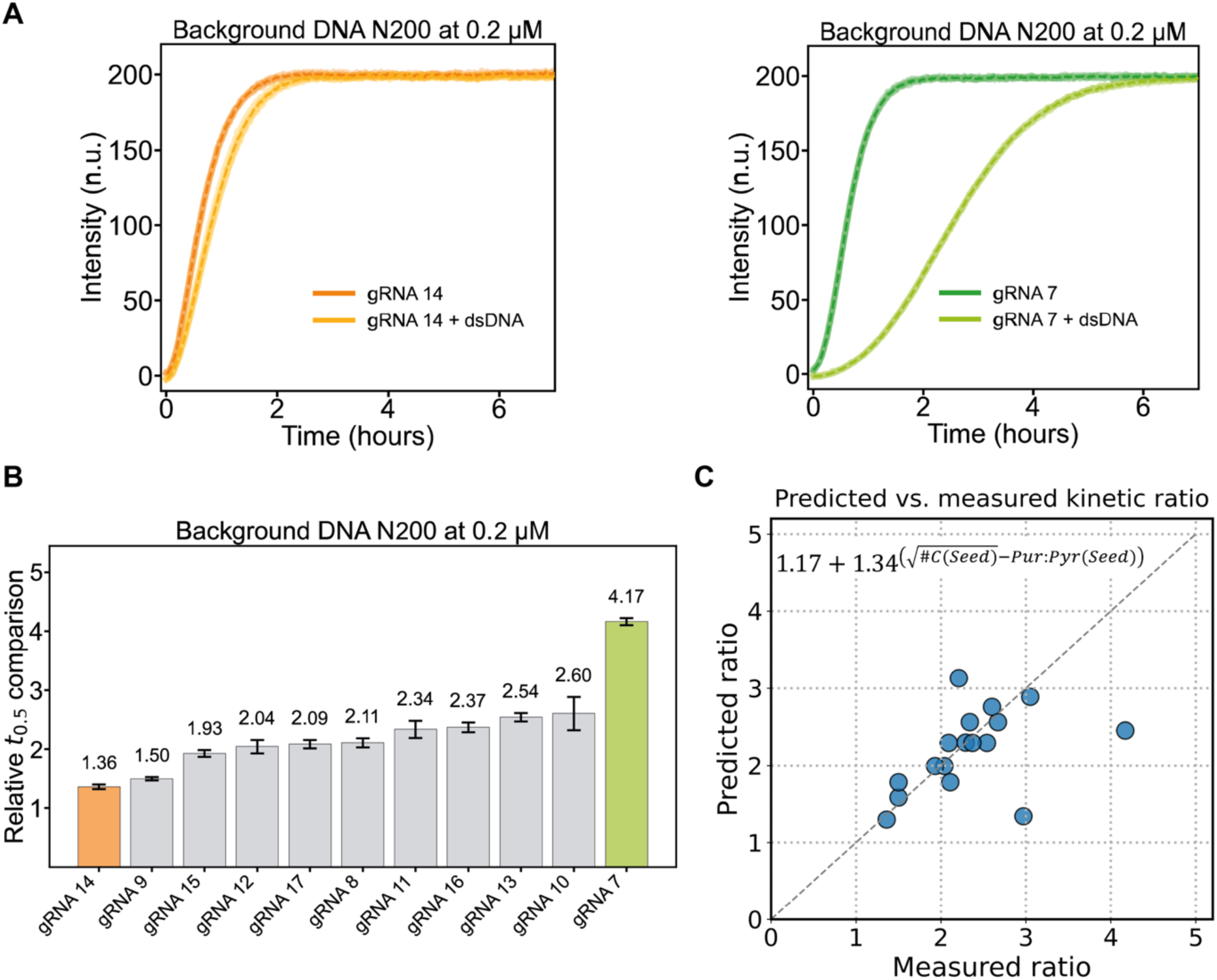
Data sets wtith varying Pur:Pyr ratio. (**A**) Fluorescence curves for the best performing gRNA 14 (left) and the worst performing gRNA 7 (right). (**B**) Bar plots showing the relative slowdown of the different gRNA versions. (**C**) Symbolic regression analysis of the data. The resulting expression given in the inset captures the trend that sequences with higher Pur:Pyr and numbers of cytosines in the seed result in less kinetic slowdown.

### Variation of Cas12a:gRNA activation *in cellulo*

We next investigated whether the sequence-dependent kinetic inhibition of Cas12a:gRNA interactions observed in vitro would also occur in cellulo. Since CRISPR/Cas activation and gene editing take place in the nucleus, both endogenous nuclear RNA and DNA are likely to influence these processes. However, the availability and reactivity of nucleic acids in the nucleus may differ substantially from in vitro conditions due to factors such as chromatin organization, molecular crowding, and other nuclear-specific effects.

Because in cellulo experiments cannot be performed without a complex nucleic acid background, we compared the CRISPR activity of two different gRNAs within the same cellular context. Specifically, we selected gRNA 14 and gRNA 7, which in our in vitro experiments (see Figure 4A) represented the least and most inhibited cases, respectively.

To assess activity, we used a CRISPR activation system in which a denAsCas12a protein fused to a VPR transcriptional activator was directed to a GFP reporter gene. As shown in Figure 5, 27 hours after transfection, cells treated with gRNA 14 and its corresponding target plasmid exhibited significantly higher GFP fluorescence than those transfected with gRNA 7.

**Figure 5.**
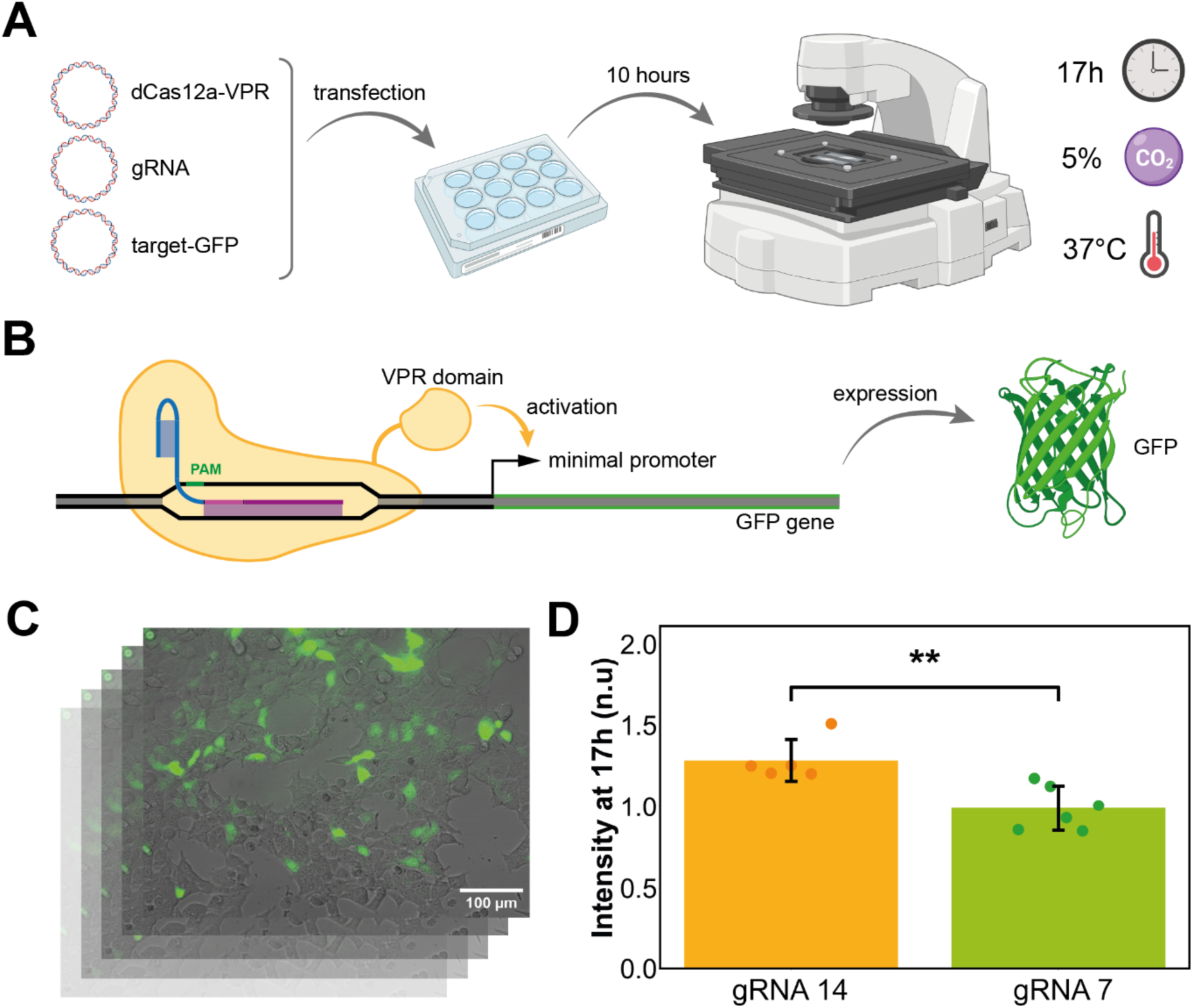
In cellulo experiment. (**A**) Workflow of the experiment. HEK293T cells are transfected with plasmids coding for denAsCas12a, a guide RNA and GFP under a minimal promoter activated by CRISPR activation upon binding of the Cas/gRNA complex to the target sequence in front of the promoter. After a 10-hour incubation the cells are placed in the incubation chamber of an EVOS microscope and images (brightfield and GFP channel) are acquired in 10-minute intervals for 17 hours. (**B**) Scheme of the CRISPR activation assay producing GFP fluorescence signal as readout. The GFP gene is under control of a minimal promoter that is only active upon activation by the VP64-p65-Rta (VPR) domain. The VPR domain is linked to a catalytically inactive (dead) enAsCas12a protein. In front of the minimal promoter there are seven targets for the used gRNA (only one shown here) which bring the modified Cas protein close enough for activation upon binding. (**C**) The images are stacked to a time-series and evaluated as described in the methods section. (**D**) Relative fluorescence intensity 27 hours post-transfection. Bars show the average intensities for cells transfected with the gRNA 14 and the gRNA 7 gRNA/target constructs. Data are acquired from 5 and 6 field of views respectively, shown by individual dots, originating from three biological replicates. Standard deviation of the mean is shown as error bar for each condition. gRNA 14 samples show higher fluorescence signal than the gRNA 7 samples. The difference between both conditions is significant with p = 0.00433. Significance was tested with a Mann-Whitney-U test based on rejection of normal distribution by a Shapiro-Wilk test and Levene’s test confirmation of equality of variances.

This outcome is consistent with the in vitro findings. Although only two gRNA sequences were tested in this in cellulo setting, the results support the idea that gRNA 14’s reduced susceptibility to background interactions improves both the kinetics and efficiency of CRISPR-mediated gene activation.

## DISCUSSION

### Main findings

The selection of target sequences and the design of guide RNAs are critical steps in the implementation of CRISPR/Cas technologies, enabling diverse applications ranging from biosensing and biocomputing to gene and cell therapies. In cellular environments, Cas-gRNA complexes do not encounter isolated target DNA strands but must operate within a dense molecular milieu comprising numerous non-cognate DNA and RNA molecules - the genome and transcriptome, respectively. Similarly, in in vitro biosensing applications, specific target sequences must be reliably detected within the complex background of clinical samples. While significant efforts have been made in recent years to improve gRNA design to minimize off-target activity in vivo and reduce false positives in biosensing, less attention has been given to the kinetic variability that may arise from such off-target interactions.

In the present work, we systematically investigated how various factors influence the kinetics of Cas12a:gRNA target binding in complex sequence environments. Using a bottom-up in vitro approach, we first examined the kinetics of target search in the presence of designed background sequence pools, which allowed us to correlate the observed kinetic inhibition with specific design features, such as the presence of PAM motifs, seed region complementarity, or combinations thereof (Figure 2). We next investigated the susceptibility of different gRNA sequence designs to interference from DNA and RNA strands with randomized sequences.

This set of experiments suggested a strong influence of the GC content and the purine-to-pyrimidine ratio within the target recognition site (spacer) of the gRNA (Figure 3 & 4). Across 17 tested gRNAs with varying sequence features, the most favorable performance was observed for gRNAs with high AU:GC ratio and simultaneously high purine content, while sequences rich in cytosines (i.e., high GC and high pyrimidine content) showed the strongest kinetic inhibition by background dsDNA sequences. The overall trend can be well captured in a simple expression derived with a symbolic regressor.

Initial experiments in human cells indicate that the kinetic effects observed in vitro may also influence CRISPR activity in a cellular context. We monitored GFP expression following CRISPR activation in HEK293T cells using a catalytically inactive Cas12a variant. Although this version does not cleave DNA, it still undergoes target search and is subject to the same background interactions with endogenous RNA and genomic DNA. Among the two tested gRNA designs, the high Pur:Pyr variant (gRNA 14) resulted in stronger gene activation than the one with a low Pur:Pyr ratio (gRNA 7). While this comparison is limited to two representative sequences, the results support the notion that background-induced kinetic modulation also occurs in vivo. A more systematic analysis will be required to fully understand these effects in cellular environments and biosensing applications.

### Interpretation

To interpret our findings we have to distinguish the different effects of RNA and DNA backgrounds as well as thermodynamic and kinetic effects:

-*RNA background*: Our results indicate that Cas12a activity is only weakly affected by the presence of an RNA sequence background, as a 200-fold excess relative to the target DNA concentration was required to observe a measurable effect. Two types of effects must be considered in this context. First, unbound guide RNAs may hybridize with random RNA sequences to varying degrees, potentially forming inhibitory secondary structures that prevent loading into the Cas12a protein. Second, once the guide RNA is incorporated into the Cas12a complex, it may still interact with surrounding RNA molecules. However, since Cas12a does not naturally target RNA, we consider such interactions to play only a minor role. Previous experiments with antisense RNAs for Cas9 gRNAs suggest that their presence reduces the availability of free guide RNA for Cas9 loading (38,39). Thus, the primary effect of an RNA background is likely a reduction in the effective concentration of free guide RNAs due to nonspecific sequestration. Among the different gRNAs tested those with a high purine content were least affected by the RNA background and the three-letter alphabet 3L-A (gRNA 3) sequence (only containing G,C,U) was most affected. This increased susceptibility potentially arises from the reduced sequence diversity combined with the high potential for both canonical (G–C, C–G, U-A) and wobble (G–U, U–G) base pairing, which enhances the probability of spurious interactions with background RNAs.

-*DNA background*: The effect of a double-stranded DNA background is much more pronounced. Consistent with previous findings, the strongest interactions were observed when the Cas12a:gRNA complex encountered background sequences containing both a matching PAM and an adjacent seed region. In contrast, background sequences with only a PAM but no matching seed, or with a seed region but no PAM, did not lead to appreciable interactions.

We can estimate the number of possible interaction sites for the Cas12a:gRNA within our 200 nM random dsDNA pool. Cas12a recognizes the PAM sequence TTTV, where V stands for any nucleotide except thymine (i.e., A, C, or G). The probability of finding this specific tetranucleotide in a random sequence of four nucleotides thus is: 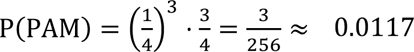 In a DNA fragment of length L = 168 base pairs, the number of possible PAM positions is 2 ⋅ (L - 4 + 1) = 330. Therefore, we expect on average N(PAM) ≈ 3.86 PAMs on one representative of the random pool, corresponding to a total PAM concentration of 772 nM in the pool. In a similar way, we can calculate the average number of sequences that contain a PAM with a fitting seed region of length m nucleotides next to it, which gives: 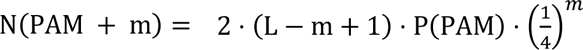 occurrences per DNA molecule. The number of occurrences exponentially drops with m – for example, the concentration of strands that contain a PAM and a neighboring 8 nt seed is only c(PAM + m) ≈ 11 pM. The observed inhibitory effect is therefore kinetic in nature, as has previously been discussed for Cas9 target search using different experimental and theoretical approaches (14,18–20,40). Interestingly, the observed kinetics of our fluorescence assay can be accurately described by a simple model in which the entire random dsDNA pool is replaced by an effective “average” sequence. This average sequence is assumed to be present at the same total concentration as the pool and to interact with Cas12a:gRNA complexes for a characteristic residence time.

In the absence of a background, we can model the rise of the fluorescence signal based on the following reactions (Model 1):

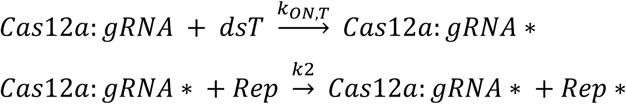

Here, dsT represents the cognate dsDNA target, Cas12-gRNA* stands for the activated effector complex, and Rep* is the fluorescent cleavage product of the doubly labeled reporter Rep. The fluorescence signal is assumed to be proportional to the concentration [Rep*]. Consistent with previous studies, we assume irreversible binding of the effector to its target (20,40) and thus disregard the unbinding reaction. In the presence of a random pool, these processes are augmented by the interaction with the background dsB (Model 2):

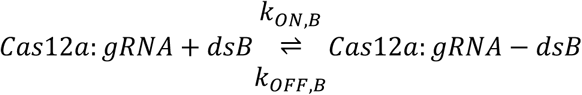

As shown in Figure 6, the models accurately reproduce the observed fluorescence time courses both in the absence and presence of the background pool. Notably, the extracted on-rates for the target and background strands differ by approximately two orders of magnitude (e.g k_ON,T_ = 1.8 × 10^4^*M*^−1^*s*^−1^, *k*_ON,B_ = 1.1 × 10^6^*M*^−1^*s*^−1^). It is important to note that k_ON,T_represents a lumped rate constant, which encompasses not only initial binding, but also subsequent conformational changes and activation of the nuclease domain within the Cas12a:gRNA effector complex.

**Figure 6.**
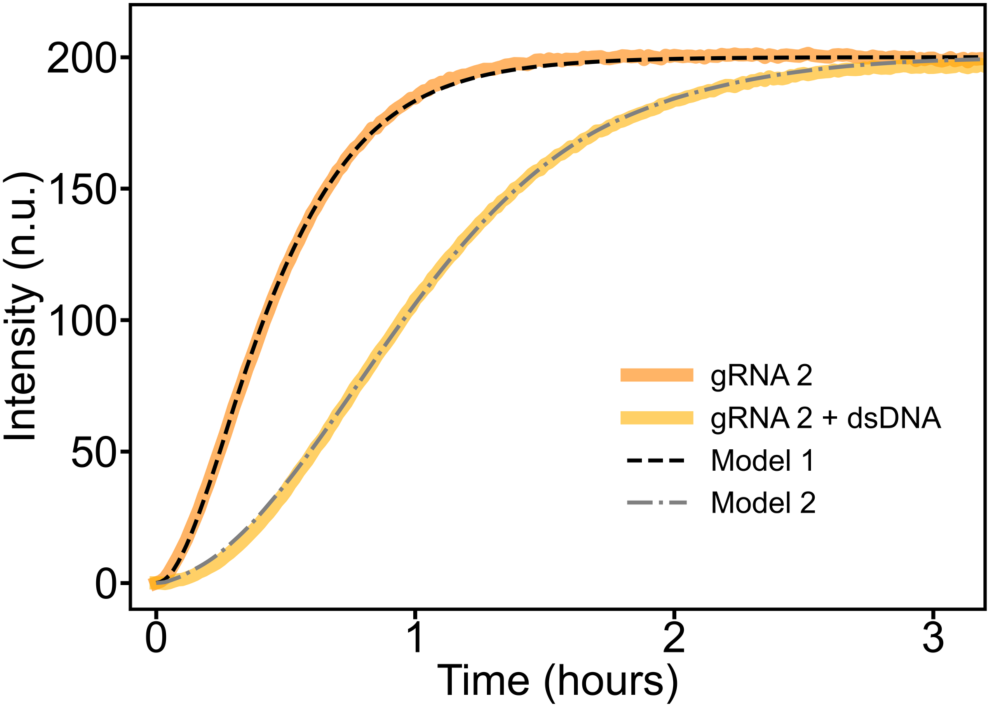
Simple kinetic model for the time course of the DETECTR reaction. Shown is the experimentally determined time-course for gRNA 2 and its cognate target in the absence and presence of a random DNA pool. Dashed and dash-dotted lines show the results of a simulation with the simple kinetic Model 1 and 2 as explained in the text.

In the context of this model, the interaction of effector complexes with gRNAs of different sequence composition with the sequence background is reflected in different unbinding rates *k*_OFF,B_. In the example shown in Figure 6, the off rate is *k*_OFF,B_ = 0.026 *s*^−1^. This corresponds to an effective equilibrium dissociation constant of *K*_*D,eff*_ = k_OFF,B_/k_ON,B_ = 24 nM. This is similar to the K_D_ for a 10 bp long duplex.

-*Sequence dependence*: Why are gRNAs with high purine-to-pyrimidine ratios in the seed less affected by a dsDNA background? Because we quantified the kinetic ratio r (the slowdown caused by the random dsDNA pool), the observation could reflect (i) faster or more favorable binding/activation on the cognate target, (ii) reduced off-pathway interactions with non-cognate background DNA, or (iii) a combination of both. In hybrid strand-displacement experiments with RNA invading DNA duplexes, a high purine content in the invading RNA strand has been reported to accelerate displacement of the DNA (29), a similar effect may operate during Cas complex engagement with its DNA target. Moreover, higher A/U content and purine enrichment can reduce intra-gRNA secondary structure, potentially yielding a more accessible seed region that accelerates R-loop formation at the cognate site. Conversely, AU-rich seeds form less stable partial or mismatched heteroduplexes with random background DNA, and thus dissociate more rapidly from non-cognate sites, while still nucleating efficiently at the true target where full complementarity stabilizes the complex.

We were able to derive a simple expression using symbolic regression that captures the trend in r values for 14 out of 17 tested gRNAs. This expression suggests that a high purine-to-pyrimidine ratio and a low number of cytosines in the seed region are favorable. However, three of the 17 gRNAs do not conform to this trend, indicating that more complex behaviors are involved. For instance, our most strongly affected gRNA, gRNA 7, has a large number of uracils in the seed, which may relate to the formation of hybrid wobble base pairs with background DNA. Remarkably, our fastest gRNA, gRNA 14 (r = 1.36), has a seed sequence very similar to that of gRNA 6, which nevertheless exhibits a kinetic ratio r of 3.

Overall, the sequence dependence thus appears to be complex. From a practical perspective, however, we believe that the derived expression may serve as a useful rule of thumb for gRNA design—bearing in mind that it does not capture the full complexity of the system.

-*In vivo relevance:* Our initial experiments suggest that designing guide sequences with a specific GC content and purine-to-pyrimidine ratio can be beneficial for target search and recognition, even in vivo. However, the effect is considerably less pronounced than under in vitro conditions. This is not surprising, given that the intracellular environment is far more complex due to factors such as compartmentalization, varying concentrations, and the different accessibility of binding partners. Within the nucleus, for example, the DNA concentration is on the order of micromolar, but only a fraction of the genomic DNA is accessible due to chromatin compaction. It has been noted that chromatin structure plays a crucial role in modulating target recognition by CRISPR–Cas effectors (41). In addition, a potentially interfering RNA background is present in the nucleus, consisting of abundant mRNAs as well as various regulatory RNA species such as long non-coding RNAs, snoRNAs, and others, many of which localize to specific membraneless subcompartments. Such factors, along with those investigated in this study, interact in complex ways and must be considered collectively to achieve a holistic understanding of CRISPR–Cas target search and binding kinetics in cellular environments.

## DATA AVAILABILITY

Python scripts, normalized data and a list of sequences are available at https://github.com/TomTom9595/Cas12a-Complex-Environment.

## AUTHOR CONTRIBUTIONS

Thomas Mayer: Data curation, formal analysis, investigation, visualization, writing - original draft. Henning Hellmer: Data curation, methodology, investigation, visualization, writing - original draft. Friedrich C. Simmel: Conceptualization, project administration, validation, simulation, funding acquisition, writing - original draft.

## ACKNOWLEDGEMENTS

The authors thank Jara Meier and Alexander Hess for support with lab work and Lukas Oesinghaus for helpful discussions. Parts of Figure 5 were created with BioRender.com, the publication license can be found in the GitHub repository.

## FUNDING

This work was supported by the Deutsche Forschungsgemeinschaft through DFG grant SI761/5−1 (DFG project number 453249455) and CRC392 TP A5 (DFG project number 521256690 – TPA5).

## CONFLICT OF INTEREST

None declared.

## Notes

### Competing Interest Statement

The authors have declared no competing interest.

